# Deep vein thrombosis-primed neutrophils reshape the brain microenvironment to promote brain metastasis

**DOI:** 10.64898/2026.09.15.751589

**Authors:** Shun Endo, Denys Rujchanarong, Karen Keeran, Kenneth Jeffries, Wei Zhang, Maxwell Bannister, Keita Saeki, Debbie Wei, Andy Tran, Kate Ellison, Anna Lee Fong, Langston Lim, Michael Kruhlak, Ross Lake, Christian A. Combs, Patricia Steeg, Michael Kelly, Stanley Lipkowitz, Yogendra Kanthi, Takeo Fujii

**Affiliations:** Women’s Malignancies Branch, Center for Cancer Research, National Cancer Institute, National Institutes of Health, Bethesda, MD 20892, USA; Laboratory of Cancer Biology and Genetics, Center for Cancer Research, National Cancer Institute, National Institutes of Health, Bethesda, MD 20892, USA; Animal Surgery and Resources Core Facility, National Heart, Lung, and Blood Institute, National Institutes of Health, Bethesda, MD 20892, USA; Medical Scientist Training Program, Johns Hopkins University School of Medicine, Baltimore, MD 21205, USA; Section on Molecular Genetics of Immunity, Eunice Kennedy Shriver National Institute of Child Health and Human Development, National Institutes of Health, Bethesda, MD 20892, USA; CCR Microscopy Core, Center for Cancer Research, National Cancer Institute, National Institutes of Health, Bethesda, MD 20892, USA; Single Cell and Spatial Core, Cancer Research Technology Program, Frederick National Laboratory for Cancer Research, Frederick, MD 20701, USA; LCBG Microscopy Core, Laboratory of Cancer Biology and Genetics, Center for Cancer Research, National Cancer Institute, National Institutes of Health, Bethesda, MD 20892, USA; Light Microscopy Core, National Heart, Lung, and Blood Institute, National Institutes of Health, Bethesda, MD 20892, USA; Section of Vascular Thrombosis & Inflammation, Division of Intramural Research, National Heart, Lung, and Blood Institute, National Institutes of Health, Bethesda, MD 20892, USA

## Abstract

Brain metastasis (BrM) is a devastating complication of triple-negative breast cancer (TNBC), yet how host macroenvironmental conditions influence brain metastatic susceptibility remains poorly understood. Here, we investigated the impact of deep vein thrombosis (DVT), a common complication in patients with cancer, on TNBC brain metastasis. Analysis of a large clinical cohort identified DVT as an independent risk factor for BrM in patients with metastatic breast cancer. Using two syngeneic TNBC models, we demonstrated that DVT selectively enhanced brain metastasis without significantly affecting primary tumor growth or lung metastasis. Mechanistically, DVT increased neutrophil accumulation in the brain metastatic microenvironment, and neutrophil depletion completely abrogated the metastasis-promoting effect of DVT. Single-cell RNA sequencing of CD45+ cells in peripheral blood revealed that DVT reprogrammed circulating neutrophils toward migratory and inflammatory transcriptomic profiles characterized by neutrophil extracellular trap (NET) formation. DVT increased CXCR2 expression on circulating neutrophils and enhanced neutrophil recruitment in brain. Moreover, DVT significantly increased circulating NETs. Collectively, our findings identify DVT as a systemic driver of TNBC brain metastasis and reveal that DVT primes circulating neutrophils toward enhanced vascular recruitment and NET formation, providing a mechanistic link between cancer-associated thrombosis and brain metastatic susceptibility.

## Introduction

Among distant organ cancer metastases, brain metastasis (BrM) is the most devastating with significant impairment of neurological function, quality of life, and overall prognosis^1^. Particularly, BrM in triple-negative breast cancer (TNBC) has an extremely poor prognosis with the median overall survival of four to five months^2^. Approximately 25-50% of patients with metastatic TNBC develop BrM^3–5^. Despite significant advances in the systemic therapies for extracranial diseases, treatment options for TNBC BrM remain limited and are primarily radiation or surgical resection with limited efficacy^6^. Therefore, developing novel interventional strategies through the mechanisms that make the brain vulnerable to metastasis is an unmet clinical need.

It is increasingly recognized that the host macroenvironment, comprising systemic host factors, plays a central role in metastatic niche formation^7^. Particularly, inflammation may create a pro-metastatic niche in distant organs ^8–10^. Despite the essential roles of inflammation in cancer progression, non-specific anti-inflammatory agents have failed in clinical trials ^11,12^, highlighting the need for deeper mechanistic understanding of systemic inflammation milieu that facilitate cancer metastasis.

Deep vein thrombosis (DVT) is a common complication among patients with cancer, induces systemic pathophysiological changes, and is associated with poor survival outcomes^13–15^. Our prior work identified that DVT is associated with high incidence of brain metastasis in patients with metastatic breast cancer^16^. Despite the clinical data suggesting the potential impact of DVT on brain metastasis, how DVT-induced systemic macroenvironmental alterations induce neurovascular and neuroimmune remodeling resulting in the regulation of brain metastatic susceptibility is poorly understood.

Here, we show that DVT-primed circulating neutrophils are essential for the promotion of TNBC BrM. DVT-primed neutrophils obtain enhanced migratory capacity represented by upregulation of CXCR2 expression on circulating neutrophils and enhanced neutrophil extracellular traps (NETs) formation.

## Results

### DVT is an independent risk factor for brain metastasis in breast cancer

In our previous retrospective study of 78 patients with *de novo* stage IV breast cancer, we identified venous thromboembolism (VTE), including DVT and pulmonary embolism (PE), as a significant clinical risk factor for the development of brain metastasis^16^. To validate this association in an independent, large-scale cohort, we analyzed data from the TriNetX Research Network^17^. Patients with extracranial metastatic breast cancer were identified and stratified according to the presence or absence of DVT (Supplementary Fig. 1A).

Because patients with more advanced and extensive disease are at increased risk of both DVT and brain metastasis, propensity score matching was performed to minimize baseline differences of tumor metastatic profile between groups. After matching, 2,511 patients were included in each cohort (Supplementary Table 1). To reduce potential reverse causality, brain metastases occurring within one year of the diagnosis of metastatic disease were excluded, and follow-up for incident brain metastasis began one year after diagnosis. The final analysis included 2,226 patients in the DVT cohort and 2,326 patients in the non-DVT cohort. The incidence of the brain metastasis was significantly higher in the DVT group (5.03% vs. 2.79%, Odds ratio 1.8, P-value=9×10⁻⁵, Supplementary Table 2). Patients with DVT exhibited a significantly higher cumulative incidence of brain metastasis over follow-up compared with patients without DVT (HR 1.67, 95% CI 1.2–2.3, P = 9.5×10⁻¹⁵, Fig 1A). Here, we found again that DVT is an independent risk factor for brain metastasis in matched cohorts of patients with extracranial metastatic breast cancer.

**Figure 1.**
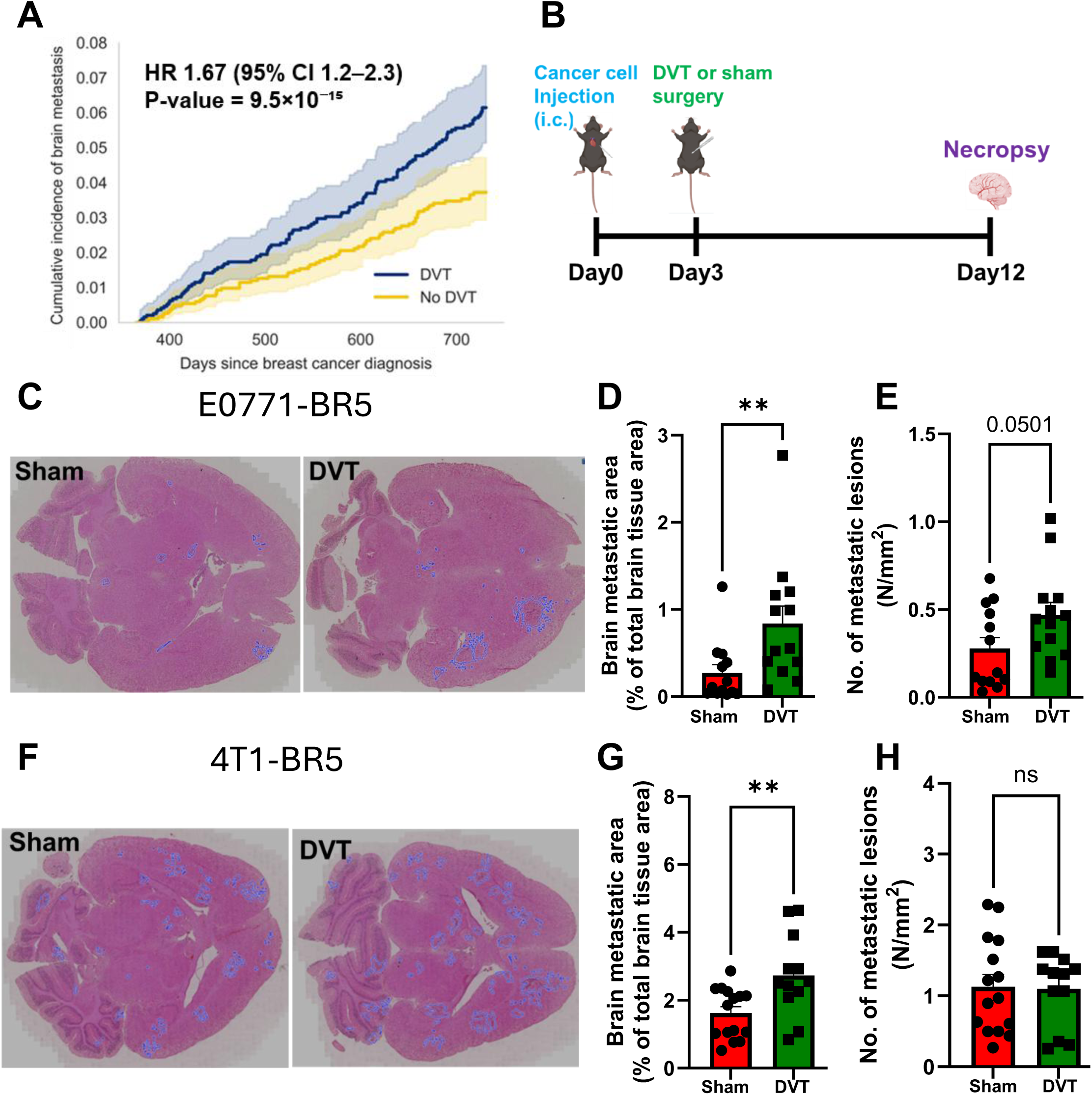
DVT promotes brain metastasis. (A) Cumulative incidence of brain metastasis in patients with metastatic breast cancer complicated by DVT (blue) or no (yellow). (B) Schematic of animal model recapitulating patients with metastatic breast cancer complicated with DVT. (C-E) E0771-BR5 model, H&E staining of brain sections at endpoint (Day 12) (C) Brain metastatic area (D), number of brain metastatic lesions (E) (n=13 for sham, n=13 for DVT, three independent experiments). (F-H) 4T1-BR5 model, H&E staining of brain sections at endpoint (Day 12) (F) Brain metastatic area (G), number of brain metastatic lesions (H) (n=15 for sham, n=12 for DVT, four independent experiments).

### DVT selectively promotes TNBC brain metastasis in immunocompetent animal models

To determine the effects of DVT on TNBC brain metastasis, we developed an animal model simulating the patients with disseminated cancer cells complicated with DVT. DVT was induced using a well-established clinically relevant Electrolytic Inferior Vena Cava Model (EIM)^18^. The DVT or sham surgeries were performed 3 days after intracardiac injection of syngeneic brain tropic cancer cells to test the effects of DVT on early colonization and outgrowth (Fig. 1B). Representative images of the thrombus at the endpoint (Day 12) are shown in the Supplementary Fig. 1B and 1C. The immunofluorescent staining demonstrated neutrophil infiltration in the thrombi as reported previously^19^ (Supplementary Fig. 1D). DVT increased brain metastasis area 1.5- to 3-fold (E0771-BR5; Fig 1C-D, 4T1-BR5; Fig. 1F-G), supporting that DVT has a critical role in early metastatic colonization. There was a trend toward an increased number of metastatic lesions following DVT in the E0771-BR5 (Fig. 1E), but not in the 4T1-BR5 (Fig. 1H), suggesting the effects of DVT on metastatic seeding or lesion initiation may be tumor type-dependent. There was no correlation between brain metastatic burden and either thrombus length or weight, indicating that the brain metastasis-promoting effect of DVT was independent of DVT severity (E0771-BR5; Supplementary Fig. 1E-F, 4T1-BR5; Supplementary Fig. 1G-H).

To determine whether the tumor promoting effect of DVT extends to primary tumor growth and metastasis to other organs or is specific to brain metastasis, parental 4T1 cells were injected into the 4^th^ mammary fat pad followed by DVT or sham surgery 3 days after orthotopic cancer cell injection (Supplementary Fig. 1I). DVT did not significantly promote primary mammary tumor growth (Supplementary Fig 1J) or lung metastasis (Supplementary Fig. 1K-L). Together, these results indicate that DVT preferentially promotes brain metastasis by enhancing the early colonization and outgrowth of TNBC cells within the brain microenvironment.

### Neutrophils mediate metastasis promoting effects of DVT

Key cellular components of the tumor microenvironment (TME) were evaluated by immunofluorescent (IF) staining^20,21^. DVT did not significantly affect microglia, astrocytes, angiogenesis, or infiltration of CD3+ T cells (Fig. 2A-D). Intriguingly, neutrophil infiltration within the TME was significantly increased in the DVT group (Fig. 2E-G), whereas circulating neutrophil counts remained unchanged (Supplementary Fig. 2A-B). Similarly, no significant differences in circulating CD3+, CD4+, or CD8+ T-cell counts were observed between the DVT and sham groups (Supplementary Fig. 2C-E). These findings indicate that the increased neutrophil accumulation in the brain TME was not attributable to an increase in circulating neutrophil counts.

**Figure 2.**
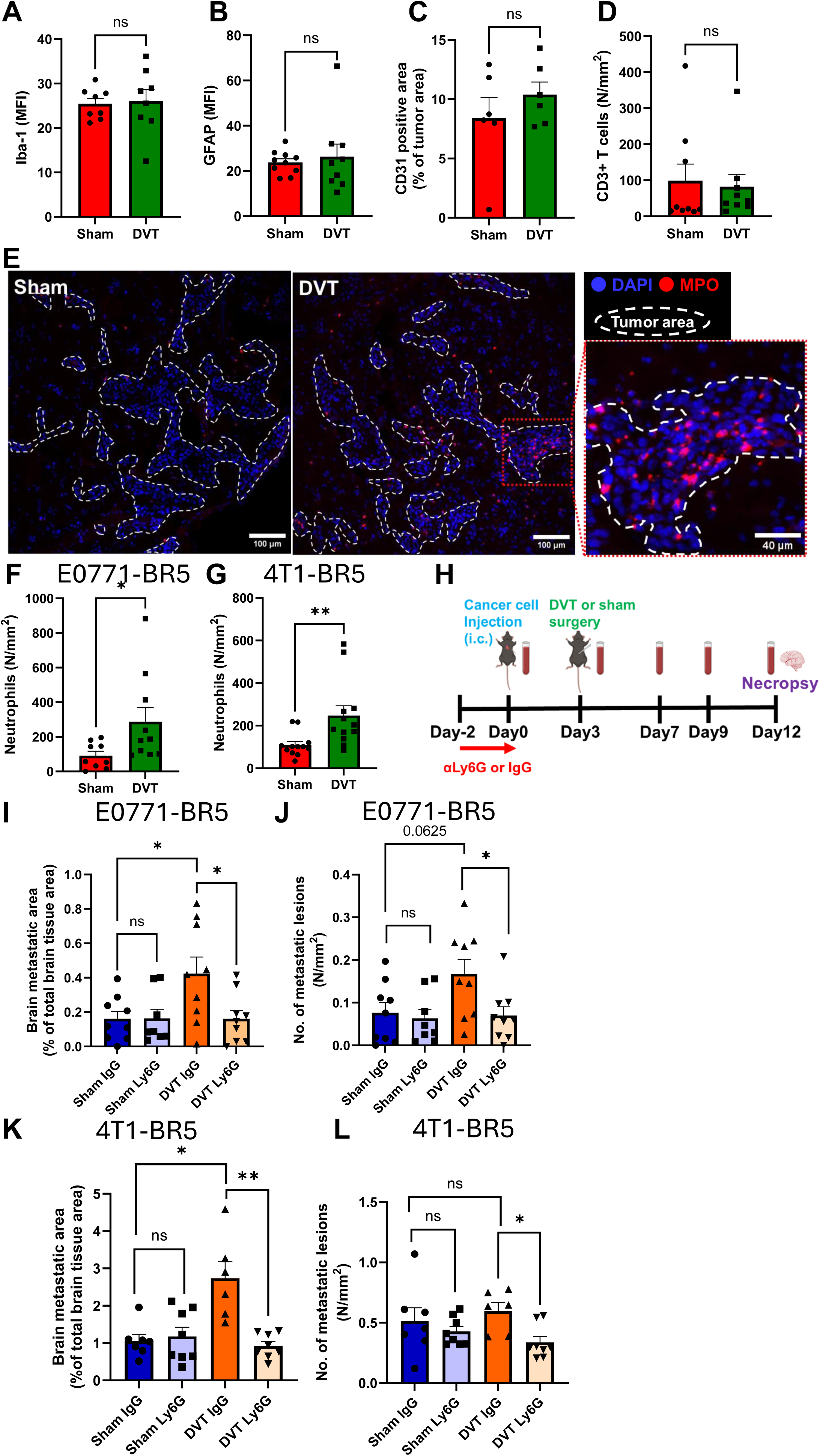
DVT-primed neutrophils regulate brain metastasis. (A-D) Quantification of immunofluorescent staining of microglia (A), astrocytes (B), Endothelial cells (C), and CD3+ T cells (D). (E) Representative images of immunofluorescent staining of MPO in brain (E0771-BR5). (F) Quantification of infiltrated neutrophils normalized to brain metastatic area in TME (E0771-BR5). (G) Quantification of infiltrated neutrophils normalized to brain metastatic area in TME (4T1-BR5). (H) Schematic of DVT-brain metastasis model with neutrophil depletion. (I) Quantification of brain metastatic area at the endpoint (Day 12, E0771-BR5, three independent experiments). (J) Quantification of the number of brain metastatic lesions at the endpoint (Day 12, E0771-BR5, three independent experiments). (K) Quantification of brain metastatic area at the endpoint (Day 12, 4T1-BR5, three independent experiments). (L) Quantification of the number of brain metastatic lesions at the endpoint (Day 12, 4T1-BR5, three independent experiments).

To understand the functional roles of neutrophils in TNBC brain metastasis, neutrophil depletion experiments were performed. Given the established role of neutrophils during the early phases of metastasis^22,23^, neutrophils were selectively depleted during the early colonization phase (Fig. 2H). The efficiency and kinetics of neutrophil depletion were assessed by flow cytometry at multiple timepoints. As intended, neutrophils were effectively depleted on Day 0, when tumor cells were intracardially injected, and remained depleted on Day 3, when DVT or sham surgery was performed with gradual recovery (Supplementary Fig. 2F, G). Intriguingly, transient neutrophil depletion during the early colonization phase completely abrogated the DVT-induced increase in brain metastatic area in both the E0771-BR5 and 4T1-BR5 models (E0771-BR5; Fig 2I, 4T1-BR5; Fig 2K). Although DVT did not significantly increase the number of metastatic lesions, neutrophil depletion reduced the number of metastatic lesions specifically in the DVT groups in both models (E0771-BR5; Fig 2J, 4T1-BR5; Fig 2L), rasing the potential role of DVT-primed neutrophils in promoting metastatic seeding. Mechanistically, DVT-promoted brain metastasis was independent of enhanced tumor cell extravasation or early proliferation (Supplementary Fig. 2H-J), indicating that DVT-primed neutrophils instead promote tumor cell survival and subsequent metastatic colonization. Together, these results demonstrate that neutrophils are critical mediators of DVT-induced enhancement of TNBC brain metastasis.

### DVT reprograms circulating neutrophils toward inflammatory and migratory phenotypes

To understand the molecular characteristics of DVT-primed circulating neutrophils, single-cell RNA sequencing was performed on FACS-sorted DAPI-negative CD45-positive cells (Fig. 3A-B). Among the four neutrophil clusters identified, the proportion of Neutrophils_2 cluster was increased in the DVT group compared with the sham group (Fig. 3C). A total of 1220 Differentially Expressed Genes (DEGs) (344 upregulated and 876 downregulated genes) were identified in the Neutrophils_2 (Fig. 3D). Genes associated with inflammation, activation, chemotaxis, and migration were enriched among the upregulated genes in this cluster (Fig. 3E). Consistent with these findings, Gene Ontology (GO) analysis demonstrated enrichment of pathways related to cell migration and chemotaxis (Fig. 3F) and KEGG analysis identified enrichment of inflammatory and activation-related pathways, including NET formation (Fig. 3G). In contrast to the neutrophil changes, no significant differences in the proportions of T cells, B cells, NK cells, or monocytes were observed between the DVT and sham groups (Fig 3A-C). Consistent with the single-cell RNA-sequencing findings, flow cytometric analysis showed no notable changes in circulating T-cell populations (Supplementary Fig. 2C-E)

**Figure 3.**
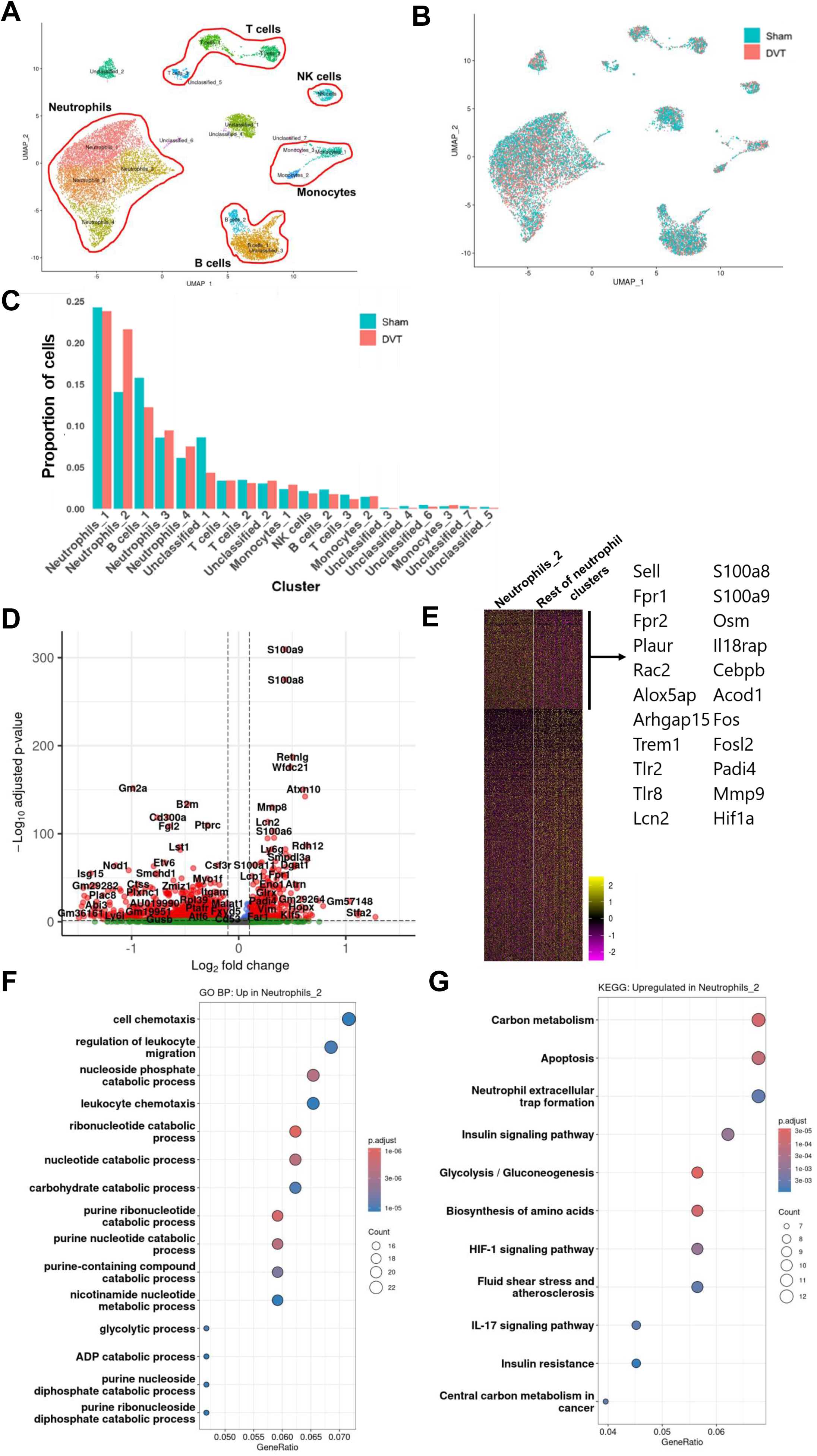
DVT-primed neutrophils are more migratory and prone to form NETs. (A) UMAP of clusters by CD45+ cells in peripheral blood (single-cell RNA sequencing, Day5). (B) UMAP comparing sham and DVT groups. (C) Proportion of cells for each cluster. (D) Volcano plot comparing the Neutrophil cluster_2 and the rest of the neutrophil clusters. (E) Heat map of relative expression of genes comparing the Neutrophil cluster_2 and the rest of the neutrophil clusters. (F) Gene Ontology (GO) term analysis of enriched pathways comparing the Neutrophil cluster_2 and the rest of the neutrophil clusters (n = 1 mouse/group). (G) KEGG pathway enrichment analysis comparing the Neutrophil cluster_2 and the rest of the neutrophil clusters (n = 1 mouse/group).

### DVT-primed neutrophils highly express CXCR2

To determine whether DVT enhances the migratory phenotype of circulating neutrophils, we examined the expression of neutrophil surface molecules involved in chemotaxis, tethering and rolling, firm adhesion, and intravascular crawling by flow cytometry^24–26^. Among eight candidate surface markers, CXCR2 and CD62L were consistently upregulated on circulating neutrophils following DVT in both the E0771-BR5 and 4T1-BR5 models (Fig. 4A. Supplementary Fig. 3A). The upregulation of CXCR2 and CD62L was also observed in tumor naïve animals (C57/B6 in Supplementary Fig. 3B, Balb/c in Supplementary Fig. 3C), indicating that DVT-induced CXCR2 and CD62L upregulation occurs independently of tumor burden. CXCR2 signaling has been implicated in neutrophil intravascular crawling and transendothelial migration^27,28^.

**Figure 4.**
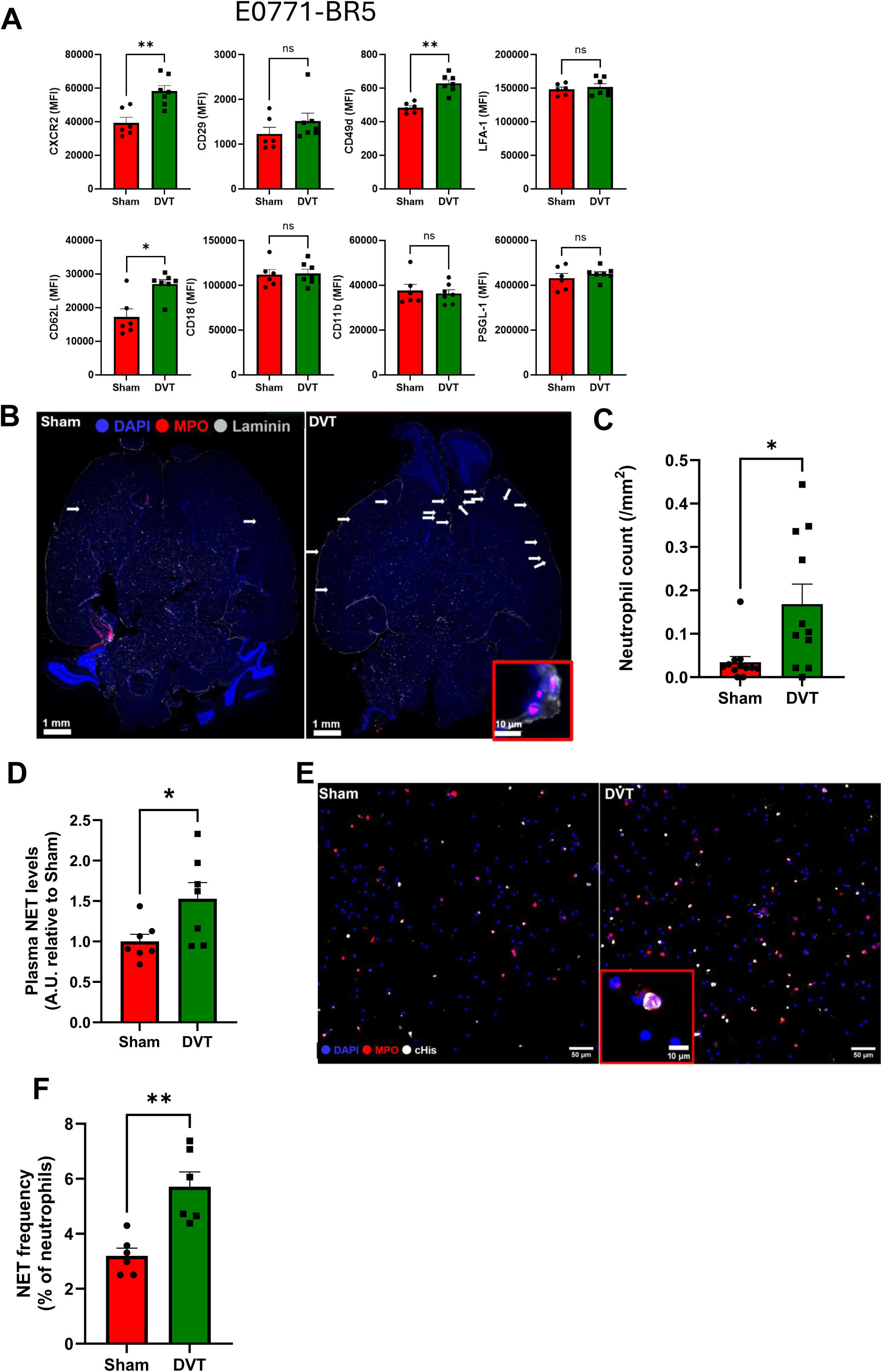
DVT-primed neutrophils upregulates CXCR2 expression and enhance NET formation. (A) Neutrophil surface markers of peripheral blood collected from E0771-BR5 injected mice (Day5) (B) ) Representative images of immunofluorescent staining of MPO and Laminin in brain (Day 5, E0771-BR5). (C) Quantification of the vasculature-associated and infiltrated neutrophils in brain (Day 5 , E0771-BR5) (D) ELISA analysis of plasma samples for NET from sham and DVT groups (Day 5). (E) Representative images of immunofluorescent staining of MPO and cHis for spontaneous NETs in peripheral blood (Day 5, E0771-BR5). (F) Quantification of the proportion of spontaneous NETs among the total number of neutrophils NETs in peripheral blood (Day 5, E0771-BR5).

Indeed, neutrophil infiltration was completely inhibited in CXCR2-deficient mice in a TNF-induced acute-inflammation model^27^, suggesting the critical role of CXCR2 in neutrophil infiltration independent of CD62L expression. Therefore, we next quantified neutrophil recruitment in the brain. Consistent with the increased CXCR2 expression, DVT significantly increased the number of neutrophils associated with the cerebral vasculature (Fig. 4B-C). Together, these findings demonstrate that DVT induces a CXCR2-high phenotype in circulating neutrophils leading to the increased recruitment of neutrophils in brain, supporting a potential role of CXCR2 in DVT-induced neutrophil recruitment to the brain.

### DVT-primed neutrophils form neutrophil extracellular traps (NETs)

Given that KEGG pathway analysis of the scRNA-seq data identified NET formation as an enriched pathway, we next investigated whether DVT promotes NET formation. NETs are extracellular DNA meshes associated with granular enzymes^29–31^ and have been implicated in multiple pro-metastatic processes, including enhanced tumor-cell migration and invasion^30,32^, chemotherapy resistance^33^, immunosuppressive niche formation^34^, suppression of anti-tumor immunity^35^, increased platelet adhesion, activation, and aggregation ^36^, and promotion of thrombosis^37,38^. To determine whether DVT increases systemic NET formation, circulating NETs were quantified by NET ELISA as previously described^9^. Circulating NET levels were significantly elevated in mice with DVT compared with sham surgery controls in the E0771-BR5 tumor-bearing model (Fig. 4D). We next assessed NET formation ex vivo by immunofluorescence staining. Consistent with the ELISA results, neutrophils isolated from mice with DVT exhibited significantly increased NET formation compared with those from control mice (Fig. 4 E-F). Together, these findings demonstrate that DVT primes circulating neutrophils toward enhanced NET formation, consistent with the NET-associated transcriptional program identified by scRNA-seq.

## Discussion

Here, we leveraged the large, publicly available, real-world dataset from the TriNetX Research Network and found that DVT was significantly associated with the increased risk of brain metastasis in patients with breast cancer. To investigate the biological link between DVT and brain metastasis, we combined the well-established clinically relevant Electrolytic Inferior Vena Cava Model (EIM) DVT model^18^ with intracardiac injection of syngeneic breast cancer cells. This models the patients with metastatic breast cancer complicated by DVT and enables investigation of how DVT influences breast cancer brain metastasis. Our findings demonstrate that DVT promotes selectively brain metastasis of TNBC. Neutrophils are the key driver of the DVT-induced brain metastasis. DVT induces a systemic inflammatory state characterized by the priming and activation of circulating inflammatory neutrophils^39,40^. DVT appears to alter neutrophil functional state rather than simply inducing systemic neutrophilia. This finding is consistent with the prior reports that inflammatory insults can activate pro-metastatic neutrophil populations that facilitate metastasis^41,42^. The major conceptual advance of this study is the identification of DVT not merely as a consequence or biomarker of advanced cancer, but as a potential modifier of metastatic susceptibility. Our findings indicate that a peripheral thrombotic event can reprogram circulating neutrophils toward a pro-metastatic phenotype and thereby influence tumor progression at a distant site.

Our findings suggest that CXCR2 on neutrophils may contribute to the recruitment of DVT-primed neutrophils to the brain metastatic niche. DVT increased surface CXCR2 expression on circulating neutrophils and was accompanied by increased neutrophil association with the cerebral vasculature. Notably, CXCR2 upregulation was also observed in tumor-naïve mice, indicating that this phenotype is induced by DVT itself. Further investigations into the mechanisms of CXCR2 upregulation are necessary. In the natural life cycles of neutrophils, high CXCR2 represents “fresh” neutrophils compared to “aged” neutrophils with low CXCR2 expression^9,43^. Additionally, another potential mechanism is the enhanced receptor recycling and shedding^44^. For translational purposes, future studies using a CXCR2 inhibitor ^45,46^ are warranted to establish the functional contribution of this axis to DVT-promoted TNBC brain metastasis.

NET formation represents another potential mechanism through which DVT-primed neutrophils may promote TNBC brain metastasis. NETs are web-like structures of DNA with associated cytotoxic enzymes released into the extracellular space^29^. NETs are known to promote migration and invasion^30,32^, induce chemotherapy resistance^33^, form immunosuppressive niche^34^, inhibit anti-cancer immunity^35^, and awaken dormant breast cancer cells^47^. Furthermore, NETs stimulate platelet adhesions and activation, resulting in the promotion of thrombosis formation^36^. NET-associated proteases, including neutrophil elastase and MMP9, can remodel the extracellular matrix and generate a microenvironment permissive for metastatic outgrowth^47^. We found that DVT increased circulating NET levels, consistent with the NET-associated transcriptional program identified by scRNA-seq. However, although our findings establish that DVT promotes NET formation, data supporting that NETs are required for the DVT-induced increase in brain metastasis remain unstudied. Future studies using complementary approaches to inhibit NET formation or promote NET degradation, including PAD4 inhibition and DNase I treatment, will be necessary to determine whether NETs function as a causal effector of DVT-primed neutrophils in promoting TNBC brain metastasis.

In this study, we found that neutrophils are required for DVT-promoting brain metastasis, but sufficiency was not tested. To prove sufficiency, further study will perform adoptive transfer of neutrophils collected from sham and DVT animals into mice without DVT. From a translational perspective, systemic neutrophil depletion is not clinically feasible. Further studies are warranted to determine whether neutrophil effector functions can be therapeutically targeted using CXCR2 inhibitors or NET-directed agents, including inhibitors of peptidylarginine deiminase 4 (PAD4), the enzyme required for NET formation^9,48,49^, DNase I, a known degrader of NET-associated DNA^30,36^, or disulfiram, an FDA-approved drug for alcohol dependence that also inhibits NET formation by blocking gasdermin D^50–52^. Further investigation is also needed to define the mechanisms by which DVT-primed neutrophils promote TNBC brain metastasis. Our findings provide insight into the stage of the metastatic cascade at which DVT-primed neutrophils may play a role in. DVT didn’t enhance cancer cell extravasation into the brain or the early proliferation of extravasated tumor cells, suggesting that DVT-primed neutrophils may instead facilitate subsequent survival and metastatic colonization. Several mechanisms could contribute to this, such as escape from innate immune surveillance^53–55^ or extracellular matrix (ECM) remodeling that establishes tumor-permissive niche^56,57^. Although our findings support a neutrophil-dominant mechanism, they do not exclude contributions from adaptive immunity or interactions of neutrophils with other cells including platelets or endothelial cells. Future studies will be warranted to determine whether DVT-primed neutrophils promote metastatic colonization through direct effects on disseminated tumor-cell survival, suppression of innate immune surveillance, remodeling of the brain metastatic niche, or a combination of these mechanisms.

Altogether, our study identifies DVT as a potential modifier of metastatic susceptibility rather than merely a consequence of advanced cancer. Our findings support a model in which DVT reprograms circulating neutrophils toward inflammatory, migratory, and NET-prone phenotypes that promote TNBC metastatic colonization in the brain. These findings establish a previously unrecognized link between peripheral thrombosis, systemic innate immune reprogramming, and brain metastatic progression, highlighting the importance of the host macroenvironment in determining organ-specific metastatic susceptibility. Importantly, targeting pathological effector functions of DVT-primed neutrophils, rather than broadly depleting neutrophils, may provide a translational strategy to mitigate the increased risk of brain metastasis associated with DVT.

## Methods

### Cell culture

Brain-tropic E0771-BR5 was cultured in RPMI (Gibco) supplemented with 10% FBS (Gibco), 100 U/mL penicillin/streptomycin (Gibco), 1% Glutamax (Gibco) in 5% CO2 atmosphere at 37 °C. Brain tropic 4T1-BR5 and parental 4T1 were cultured in DMEM (Gibco) supplemented with 10% FBS (Gibco), 100 U/ml penicillin/streptomycin (Gibco) in 5% CO2 atmosphere at 37 °C. The E0771-BR5-eGFP cell line was generated by transfecting the cells with the eGFP plasmid made by Yu Lou. In 48 hours after transduction, cells were treated with puromycin (2 μg/mL; #A11138, Gibco) for 2 weeks. Cultured cells were tested repeatedly for mycoplasma over the course of this study and remained negative for the duration of the study.

### Mice

All procedures were approved by the National Cancer Institute Animal Care and Use Committee (ACUC) and were conducted in accordance with the NIH’s Guide for the Care and Use of Laboratory Animals. Female BALB/cJ (#000651) and C57BL/6J (#000664) mice were purchased from the Jackson Laboratory. All mice were acclimated to the animal housing facility for at least 1 week before initiating experiments. Mice were housed in temperature and humidity controlled conventional facilities on a 12-hr light-dark cycle with food and water available *ad libitum*.

### Electrolytic Inferior Vena Cava Model (EIM) to model DVT

Mice were placed on heated surface (warming pad) while anesthetized. A ventral midline incision was made, and the intestines were retracted using sterile saline soaked gauze to allow for visualization of the inferior vena cava (IVC). Next, the IVC was dissected from the surrounding tissues for a distance of approximately 1 cm infrarenal. A 30-gauge silver-coated copper wire (#KY-30-1-GRN, Electrospec) with exposed copper wire at the end was placed inside a 25-gauge needle, which was inserted into the IVC and positioned against the anterior wall (where it functioned as the anode). Another copper wire was implanted subcutaneously, completing the circuit (cathode). A constant current of 250 μA was applied for 15 min supplied by a voltage-to-current converter^58^. After removal of the needle, the abdomen was closed.

In sham animals, the needle was placed into the IVC for 15 minutes without application of the current.

### Experimental brain metastasis models

4T1-BR5 (3× 10^4^ in 100uL PBS) and E0771-BR5 (6×10^4^ in 100uL PBS) were injected intracardially, which was followed by EIM DVT or sham surgery 3 days post intracardiac cancer cell injection. All the animals were euthanized within 12 days after intracardiac cancer cell injection and the brains were collected for analysis. To deplete neutrophils, anti-mouse Ly6G (200 μg/mouse, #CP129, clone 1A8, Bio X Cell) and its IgG control (200 μg/mouse, #CP190, Bio X Cell) were injected intraperitoneally every other day.

### Primary mammary fat pad tumor and spontaneous lung metastasis models

4T1 (3× 10^4^ in 10uL PBS + 10uL matrigel) was injected into the 4^th^ mammary fat pad. Tumor length and width were measured twice a week with a caliper, and tumor volume was calculated as (length x width^2^)/2. All the animals were euthanized 21 days after orthotopic cancer cell injection and the lungs were collected for analysis of metastasis.

### Quantification of metastasis

The metastasis area and the number of metastases of the brain were determined on hematoxylin & eosin (H&E)-stained slides. The slides were scanned using Axio Scan.Z1 (Zeiss) and VS200 slide scanner (Olympus). The metastasis was analyzed using QuPath v0.7.0. software. The number of metastasis was normalized by total tissue area evaluated.

### Flow cytometry analysis and FACS sorting

Flow cytometric analyses were performed using ID7000™ Spectral Cell Analyzer (SONY). The analysis was performed using FlowJo v10 (Tree Star Inc.). All analyses were conducted at the Flow Cytometry Core at the CCR/NCI. Absolute count was done using Trucount absolute counting beads (#340334, BD Biosciences) according to the manufacturer’s instructions.

Blood was drawn and placed into EDTA-coated tubes, red blood cells were lysed in ammonium–chloride– potassium (ACK) lysis buffer (#A1049201, Gibco), and blood cells were stained with fluorochrome-labeled antibodies. Immediately before flow cytometric analysis, DAPI was added to the cells.

### Immunofluorescent staining

To fixed-frozen tissue slides, antigen retrieval was done by boiling slides in Tris EDTA buffer (10 mM Tris base and 1 mM EDTA, pH 9.0) for 10 min. Then, the tissues were incubated with a blocking buffer containing PBS with 0.1% Triton X-100 and 0.2% BSA for 30-60 mins at room temperature. The tissues were incubated with the corresponding primary antibodies overnight at 4 °C. The next day, the samples were washed three times in PBS and stained with the corresponding secondary antibodies for 2 hours at room temperature. Then, the samples were washed, counterstained with DAPI and mounted with ProLong™ Gold Antifade Mountan t(#P10144, Thermo Fisher).

### NET enzyme-linked immunosorbent assay (ELISA)

NET levels in plasma were quantified as previously described^9^. Blood was collected into an EDTA coated blood collection tube. Whole blood was then centrifuged at 2,500g for 20 minutes at 4 °C. The plasma fraction was carefully collected, aliquoted, and stored at −80°C until analysis. For NET ELISA, a 96-well Enzyme ImmunoAssay/Radio Immuno-Assay (EIA/RIA) plates were coated overnight at 4°C with anti-neutrophil elastase (#sc-55549, Santa Cruz Biotechnology, 1:250) in coating buffer (15 mM Na2CO3, 35 mM NaHCO3, pH 9.6). After coating, plates were washed with PBS and blocked with 5% BSA for 2 hours at room temperature. Following blocking, 50 μL of plasma was added to each well and incubated for 2 hours at room temperature with gentle shaking, and then the wells were washed three times using washing buffer (1% BSA, 0.05% Tween20 in PBS). Next, anti-DNA-peroxidase conjugated antibody (1:50, part of the Cell Death Detection ELISA Kit, #11544675001, Sigma) in 1% BSA was added and wells were incubated for 2 hours at room temperature. The wells were washed five times using washing buffer and then 2,20-azino-bis (3-ethylbenzothiazoline-6-sulphonic acid) (ABTS) was added. Absorbance was measured at 405 nm after 10 minutes using a SpectraMax MiniMax 300 Imaging Cytometer (Molecular Devices).

### Ex vivo NET formation assay

Whole blood was collected and RBCs were lysed using ACK lysing buffer (#A1049201, Gibco). The volume equivalent to 40 μL of the original blood in serum-free RPMI medium was plated per well on poly-l-lysine-covered 8-well μ-slides (#80804, Ibidi), then left for 30 min at 37 °C in a cell culture incubator to adhere. Cells were plated in a drop of medium in the center of the well to enhance their adhesion to the central area of the well and to avoid their deposition at the edges. Cells were subsequently incubated for 12 h with 200 nM PMA or vehicle. Cells were then fixed using 4% paraformaldehyde (PFA) in PBS for 10 min; blocked and permeabilized with PBS containing 0.1% Triton X-100, 25% FBS and 5% bovine serum albumin (BSA); and stained with antibodies to citH3 (#ab281584, Abcam) and MPO (# AF3667, R&D Systems) at 1:200 dilution in blocking buffer at 4 °C overnight. Then, the cells were washed and stained with secondary antibodies: donkey anti-goat AF647 (#A21447, Invitrogen) and donkey anti-rabbit AF568 (#A10042, Invitrogen) at 1:400 and counterstained with DAPI (1:1,000) for 2 h at room temperature. Z-stack images were acquired with a Stellaris 8 FLIM (Leica) and analyzed using Fiji to identify NETs (defined as triple co-localized MPO+, citH3+ and DNA (DAPI)+ and therefore citrullinated NETs).

### Single-cell RNA sequencing

FACS sorted DAPI-CD45+ Ly6G- and DAPI-CD45+Ly6G+ cells from whole blood were mixed with the same number of cells and was submitted to the CCR Single Cell and Spatial Core for scRNA sequencing. Cell suspensions were assessed for concentration and viability on LunaFX7 automated cell counter (Logos Biosystems) prior to single cell capture on the Chromium platform using Single Cell 3′ v4 GEM-X chemistry (10x Genomics). Libraries were prepared according to 10x Genomics assay user guide and sequenced on an Illumina NovaSeq X Plus system using a recommended run parameters on a 100-cycle kit. Sequencing reads were processed using Cell Ranger v10.0.0 (10x Genomics) and aligned to the mouse reference genome (refdata-gex-GRCm39-2024-A, 10x Genomics). The resulting ambient RNA-corrected filtered feature-barcode matrix was used for downstream analysis. All the downstream analyses were performed in R v4.5.2 using Seurat. First, the datasets were subjected to quality control, and cells with mitochondrial transcripts below 5% as well as cells with detected features between 500 and 7500 were kept and were log normalized. Putative doublets were identified and removed using the scDblFinder. Gene-expression counts were log-normalized. Principal component analysis (PCA) was performed using the first 30 principal components. A shared nearest-neighbor graph was constructed using principal components of 1–20 (FindNeighbors), and cells were clustered using the FindClusters with a resolution of 0.5. Uniform Manifold Approximation and Projection (UMAP) was calculated from principal components of 1–20 using the RunUMAP. Cluster identities were assigned manually by referring to the 20 highly expressed genes in each cluster and the expression of established lineage-specific markers. Data manipulation and visualization were performed using the dplyr and ggplot2 packages, respectively. Differentially expressed genes (DEGs) between sham and DVT groups were identified using the FindMarkers in Seurat. Volcano plots were generated using the EnhancedVolcano. Heatmap was generated using the pheatmap. Gene Ontology Biological Process enrichment analysis was performed using the clusterProfiler and the mouse annotation database org.Mm.eg.db. Terms with a Benjamini–Hochberg false-discovery rate (FDR)-adjusted P value of less than 0.05 were considered significantly enriched. Kyoto Encyclopedia of Genes and Genomes (KEGG) pathway enrichment analysis was also performed using clusterProfiler with Mus musculus specified as the reference organism.

### Statistical analysis

Data are represented as mean + s.e.m unless otherwise indicated. Unpaired two-tailed t-tests or Kruskal– Wallis test was used to compare two groups, and more than two groups were compared using one-way analysis of variance (ANOVA) with multiple comparison test. All statistical analyses, except for sequencing analyses, were performed using Prism v10 (GraphPad). A P-value < 0.05 was considered statistically significant; non-significant (ns) differences are indicated in the figures.

### Analysis of TriNetX

The TriNetX Research Network was used to assess the association between DVT and breast cancer brain metastasis. Only data from U.S. healthcare organizations (HCOs) were used, representing approximately 68 million patients from 112 HCOs. Cohorts of patients were defined using the following ICD10 codes: breast cancer (C50), deep vein thrombosis (I82.4), cerebral infarction (I63), lung metastasis (C78), bone metastasis (C79.5), brain metastasis (C79.31), liver metastasis (C78.7), other metastatic sites not including lymph nodes (C79.0, C79.1, C79.2, C79.6, C79.7), hypertension (I10), dyslipidemia (E78), diabetes mellitus (E08-E13), chronic kidney disease (N18), chronic ischemic heart disease (I25), atrial fibrillation (I48), pulmonary embolism (I26), acute myocardial infarction (I21), and heart failure (I50). Patients with breast cancer were filtered to be all female, 18 years or older, and diagnosed between January 1, 2015 and January 1, 2025.

Using propensity score matching, cohorts compared in analyses were matched for age, ethnicity, race, sites of extracranial metastatic diseases, and medical comorbidities (hypertension, diabetes, dyslipidemia, smoking, obesity, atrial fibrillation, chronic kidney disease, heart failure, and stroke). All rights reserved. Use of TriNetX data or name in this publication does not imply endorsement by TriNetX or its affiliates. All analysis and conclusion are those of the authors. This retrospective analysis is exempt from informed consent. The data reviewed is a secondary analysis of existing data, does not involve intervention or interaction with human subjects, and is de-identified per the de-identification standard defined in Section §164.514(a) of the HIPAA Privacy Rule. The process by which the data is de-identified is attested to through a formal determination by a qualified expert as defined in Section §164.514(b)(1) of the HIPAA Privacy Rule. This formal determination by a qualified expert refreshed on May 2025.

## Data Availability Statement

All data associated with this study are present in the paper or the Supplementary Materials. The raw dataset generated for the single cell RNA sequencing analysis will be deposited in the NCBI Gene Expression Omnibus (GEO) and the accession number will be reported. Additional data and materials are available from the corresponding authors upon request.

## Acknowledgements

This research was supported in part by the Intramural Research Program of the National Institutes of Health (NIH). The contributions of the NIH authors are considered Works of the United States Government. The findings and conclusions presented in this paper are those of the authors and do not necessarily reflect the views of the NIH or the U.S. Department of Health and Human Services. This work utilized the computational resources of the NIH HPC Biowulf cluster (https://hpc.nih.gov). The authors thank Dr. Yu Lo for providing the eGFP plasmid.

## Author Contributions

**Conceptualization:** Y.K., T.F.

**Methodology:** S.E., D.R., K.K., K.J., W.Z., M.B., K.S., D.W., A.T., K.E., A.F.L, L.L., M.K., R.L., C.A.C., M.K., Y.K., S.L., T.F.

**Investigation:** S.E., D.R., K.K., K.J., W.Z., M.B., K.S., A.T., K.E., A.F.L, L.L., M.K., R.L., P.S., M.K., Y.K., S.L., T.F.

**Visualization:** S.E., D.R., K.K., K.J., W.Z., A.T., R.L., T.F.

**Funding acquisition:** T.F.

**Project administration:** T.F.

**Supervision:** T.F.

**Writing–original draft:** S.E., M.B., T.F.

**Writing–review and editing:** S.E., D.R., K.K., K.J., W.Z., M.B., K.S., D.W., A.T., K.E., A.F.L, L.L., M.K., R.L., C.A.C., M.K., Y.K., S.L., T.F.

**Discussion:** S.E., D.R., K.K., K.J., W.Z., M.B., K.S., D.W., A.T., K.E., A.F.L, L.L., M.K., R.L., C.A.C., P.S., M.K., Y.K., S.L., T.F.

---

**Supplementary Figure 1.**
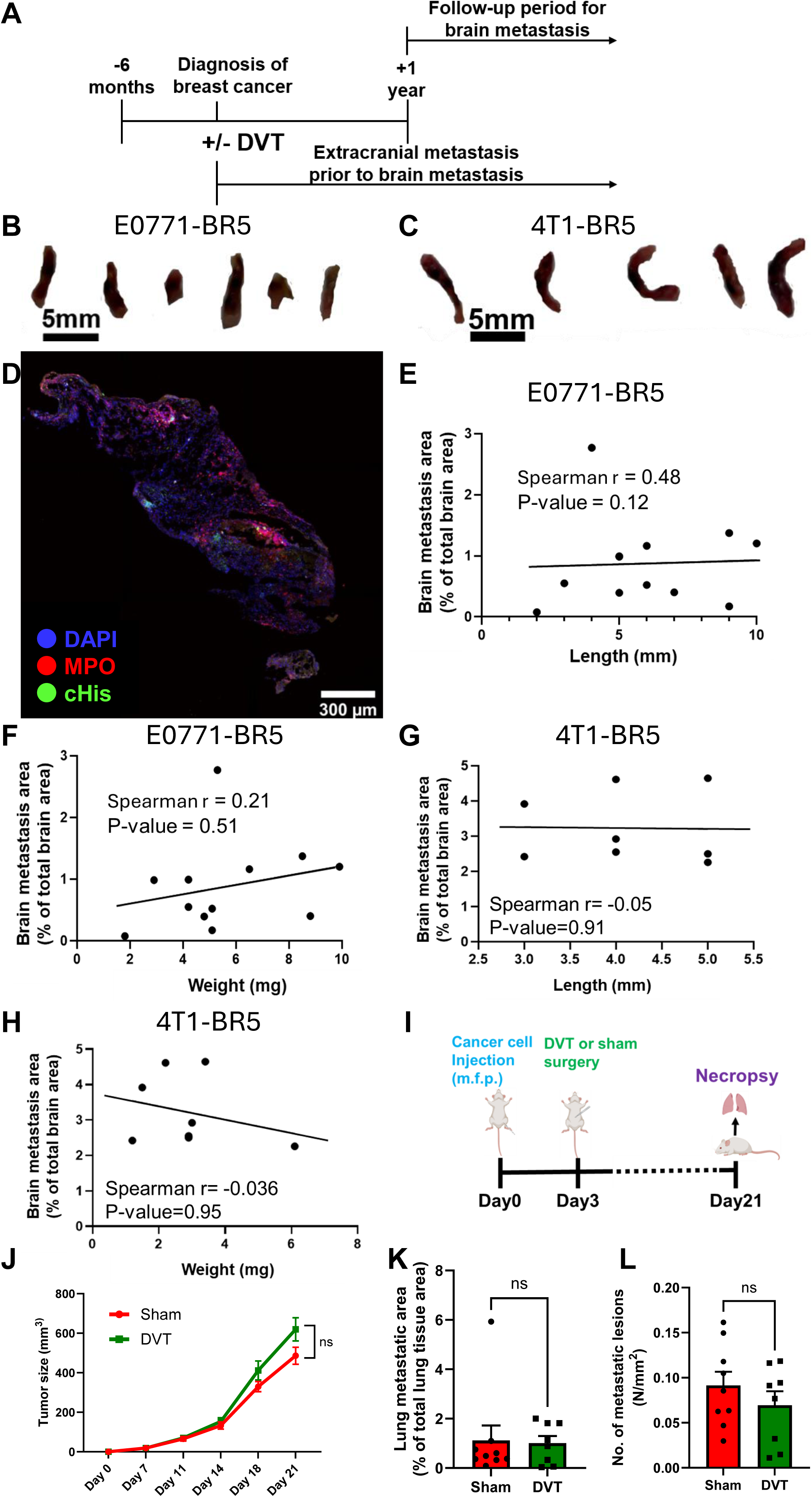
(A) Schematics of TriNetX data analysis. (B-C) Representative images of thrombus at endpoint (Day 12), E0771-BR5 (B), 4T1-BR5 (C). (D) Representative image of immunofluorescent staining of MPO and citrullinated histone (cHis) in thrombi at endpoint (Day 12) (4T1-BR5). (E-H) Spearman correlation between brain metastasis area and thrombus length and weights (E-F: E0771-BR5, G-H: 4T1-BR5) (I) Schematic of mammary fat pad injection and spontaneous lung metastasis model (4T1) (J) Tumor growth curve of mammary fat pad injection of 4T1 (n = 9 for sham, n = 8 for DVT). (K) Quantification of lug metastasis area (Day 21, (n = 9 for sham, n = 8 for DVT). (L) Quantification of the number of lug metastasis lesions (Day 21, (n = 9 for sham, n = 8 for DVT).

**Supplementary Figure 2.**
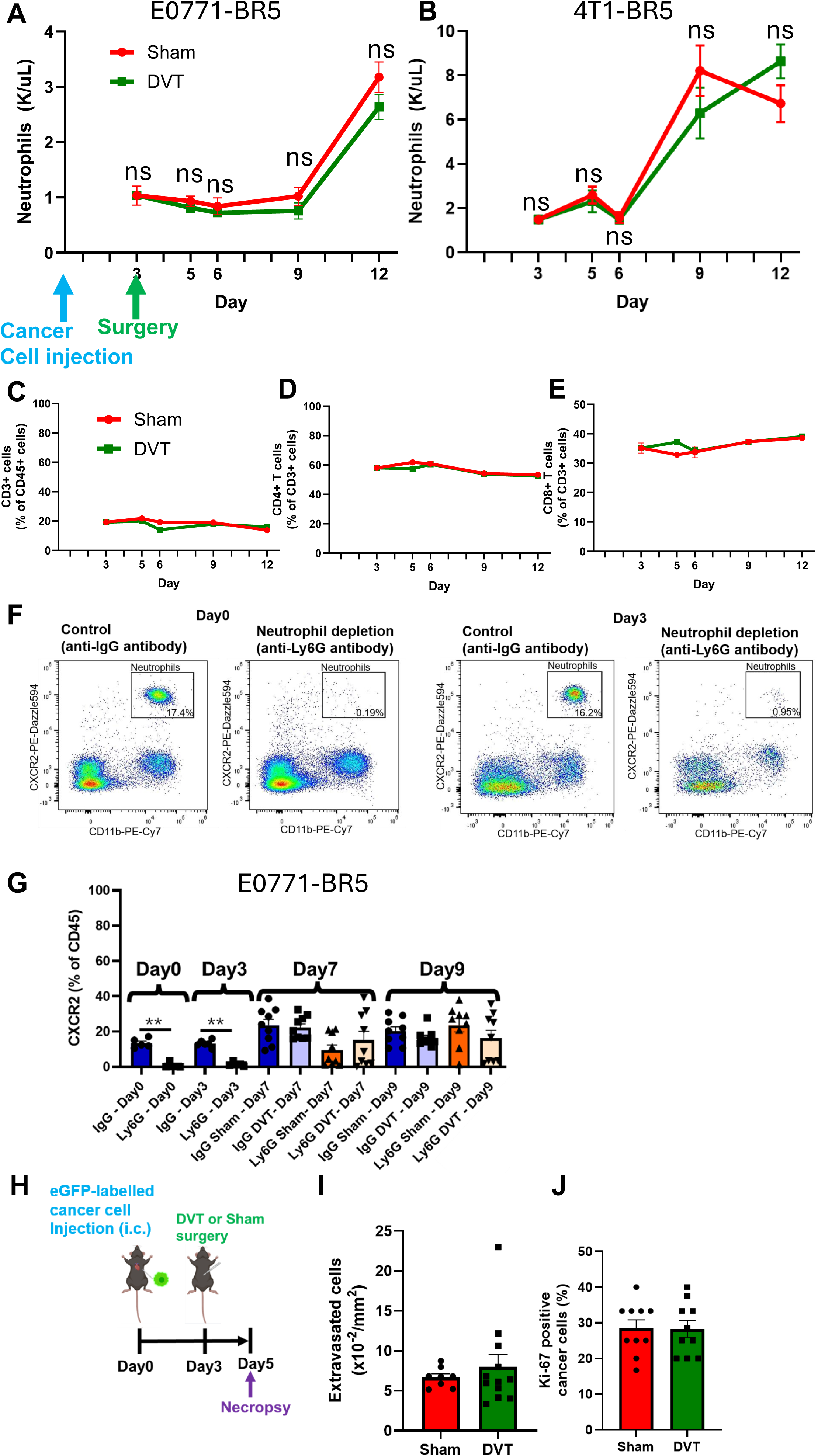
(A) Dynamics of neutrophil count in peripheral blood (E0771-BR5). (B) Dynamics of neutrophil count in peripheral blood (4T1-BR5). (C-E) Flow cytometric analysis of the proportions of CD3+ T cells (C), CD4+ T cells (D), and CD8+ T cells (E) in peripheral blood (E0771-BR5R). (F) Gating strategy of peripheral blood for neutrophils in the neutrophil depletion animal model (E0771-BR5). (G) Dynamics of neutrophil count in peripheral blood in the neutrophil depletion animal model (E0771-BR5). (H) Schematics of the DVT model with i.c. injection of eGFP-labelled E0771-BR5 cells. (I) Quantification of the number of extravasated eGFP-positive E0771-BR5 cancer cells. (J) Quantification of Ki-67 positive cancer cells among the eGFP-positive E0771-BR5 cancer cells.

**Supplementary Figure 3.**
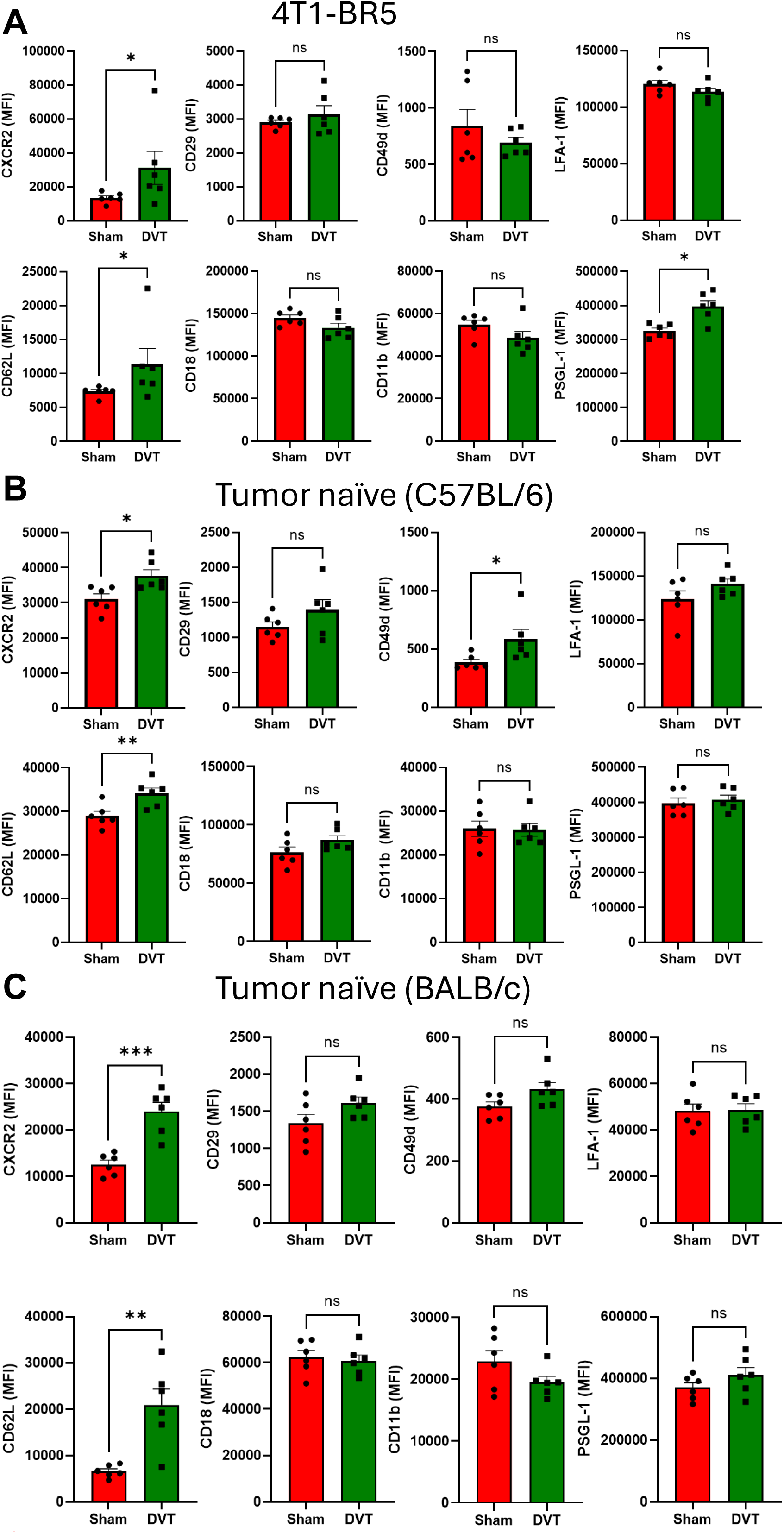
(A) Neutrophil surface markers of peripheral blood collected from 4T1-BR5 injected mice (Day5). (B) Neutrophil surface markers of peripheral blood collected from tumor naïve C57/B6 mice (2 days post surgery) (C) Neutrophil surface markers of peripheral blood collected from tumor naïve BALC/c mice (2 days post surgery).

**Supplementary Table 1.**

|  |  | **No DVT<br>(N=2,511)** | **DVT<br>(N=2,511)** |
|---|---|---|---|
| **Age at diagnosis** |  | 65 (+/-13) | 65 (+/-13) |
| **Race** |  |  |  |
|  | White | 73.84% | 72.16% |
|  | Black or African American | 11.15% | 16.29% |
|  | Asian | 6.49% | 3.74% |
|  | Native Hawaiian or Other Pacific Islander | 1.08% | 0.68% |
|  | American Indian or Alaska Native | 0.40% | 0.40% |
|  | Unknown Race | 5.34% | 3.94% |
|  | Other Race | 1.95% | 2.87% |
| **Ethnicity** |  |  |  |
|  | Not Hispanic or Latino | 84.07% | 82.36% |
|  | Hispanic or Latino | 2.55% | 5.06% |
|  | Unknown Ethnicity | 13.38% | 12.59% |
| **Estrogen receptor** |  |  |  |
|  | Positive | 37.40% | 37.32% |
|  | Negative | 14.66% | 14.38% |
| **Progesterone receptor** |  |  |  |
|  | Positive | 2.07% | 2.39% |
|  | Negative | 1.24% | 1.39% |
| **HER2 receptor** |  |  |  |
|  | Positive | 1.04% | 1.43% |
|  | Negative | 3.07% | 3.39% |
| **Initial Stage** |  |  |  |
|  | Stage 0 | 0% | 0.40% |
|  | Stage 1 | 0.64% | 0.60% |
|  | Stage 2 | 0.40% | 0.48% |
|  | Stage 3 | 0.72% | 1.00% |
|  | Stage 4 | 2.79% | 3.51% |
| **Extracranial metastasis sites** |  |  |  |
|  | Lymph nodes | 26.25% | 27.52% |
|  | Bone | 58.62% | 56.39% |
|  | Lung | 21.74% | 22.86% |
|  | Liver | 19.24% | 19.91% |
| **Medical co-morbidities** |  |  |  |
|  | Cerebral infarction | 3.54% | 3.62% |
|  | Hypertension | 47.39% | 47.59% |
|  | Dyslipidemia | 31.94% | 32.42% |
|  | Diabetes mellitus | 20.35% | 21.23% |
|  | Chronic kidney disease | 10.83% | 11.11% |
|  | Chronic ischemic heart disease | 10.75% | 11.91% |
|  | Atrial fibrillation | 6.69% | 7.77% |
|  | Acute myocardial infarction | 2.67% | 2.95% |
|  | Pulmonary embolism | 11.99% | 12.39% |
|  | Heart failure | 8.68% | 9.92% |

**Supplementary Table 2.**

|  | **No. of patients with extracranial metastasis** | **No. of patients who developed brain metastasis** | **Incidence of brain metastasis** | **Odds ratio** | **P-value** |
|---|---|---|---|---|---|
| **No DVT** | 2,326 | 65 | 2.79% | 1.8 | 9.5×10<sup>-5</sup> |
| **DVT** | 2,226 | 112 | 5.03% |  |  |

